# Understanding the Functional Properties of Lipid Heterogeneity in Pulmonary Surfactant Monolayers at the Atomistic Level

**DOI:** 10.1101/2020.07.07.191569

**Authors:** Juho Liekkinen, Berta de Santos Moreno, Riku O. Paananen, Ilpo Vattulainen, Luca Monticelli, Jorge Bernardino de la Serna, Matti Javanainen

**Affiliations:** Department of Physics, University of Helsinki, FI-00014 Helsinki, Finland; National Heart & Lung Institute, Faculty of Medicine, Imperial College London, SW7 2AZ, London, UK; Helsinki Eye Lab, Ophthalmology, University of Helsinki and Helsinki University Hospital, FI-00014 Helsinki, Finland; Computational Physics Laboratory, Tampere University, FI-33014, Tampere, Finland; MEMPHYS – Centre for Biomembrane Physics; Molecular Microbiology and Structural Biochemistry (MMSB), UMR 5086 CNRS & University of Lyon, Lyon, France; Institute of Organic Chemistry and Biochemistry of the Czech Academy of Sciences, CZ-16100 Prague 6, Czech Republic

**Author notes:** These authors contributed equally (Juho Liekkinen, Berta de Santos Moreno).

## Abstract

Pulmonary surfactant is a complex mixture of lipids and proteins lining the interior of the alveoli, and constitutes the first barrier to both oxygen and pathogens as they progress toward blood circulation. Despite decades of study, the behavior of the pulmonary surfactant is poorly understood on the molecular scale, which hinders the development of effective surfactant replacement therapies, useful in the treatment of several lung-related diseases. In this work, we combined all-atom molecular dynamics simulations, Langmuir trough measurements, and AFM imaging to study synthetic four-component lipid monolayers designed to model protein-free pulmonary surfactant. We characterized the structural and dynamic properties of the monolayers with a special focus on lateral heterogeneity. Remarkably, simulations reproduce almost quantitatively the experimental data on pressure–area isotherms and the presence of lateral heterogeneities highlighted by AFM. Quite surprisingly, the pressure–area isotherms do not show a plateau region, despite the presence of liquid-condensed nanometer–sized domains at surface pressures larger than 20 mN/m. In the simulations, the domains were small and transient, but they did not coalesce to yield a separate phase. The liquid–condensed domains were only slightly enriched in DPPC and cholesterol, and their chemical composition remained very similar to the overall composition of the monolayer membrane. Instead, they differed from liquid-expanded regions in terms of membrane thickness (in agreement with AFM data), diffusion rates, acyl chain packing, and orientation. We hypothesize that such lateral heterogeneities are crucial for lung surfactant function, as they allow both efficient packing, to achieve low surface tension, and sufficient fluidity, critical for rapid adsorption to the air–liquid interface during the breathing cycle.

## Introduction

The integrity of the alveolar gas–blood barrier is crucial for effective gas exchange and health, filtering of undesirable components, and response to inhaled hazard. At the same time, it develops tolerance mechanisms to attenuate immunopathology. The alveoli are continuously exposed to inhaled micro- and nanosized pathogens, which are normally rapidly eliminated with the help of the immune system. Immune responses in the alveoli must be tightly regulated to prevent excessive inflammation and tissue damage. Inappropriate or excessive immune responses cause the development of systemic airway inflammation, as in the Acute Respiratory Distress Syndrome (ARDS).^1^ ARDS is the major cause of respiratory failure affecting millions of people annually, and it is also a main cause of death in many viral infections, such as in severe acute respiratory syndrome coronavirus 1 (SARS-CoV-1), and in the current SARS-CoV-2 which causes the coronavirus disease 2019 (COVID-19). ^2^

The alveolar epithelium is coated by pulmonary surfactant, that is a delicate membranous proteolipid film that maintains normal lung function. Pulmonary surfactant forms a monolayer, which lines the alveolar epithelium and is synthesised and secreted by epithelial alveolar type 2 cells. Pulmonary surfactant is the most permeable interface of the human body exposed to the environment, presenting the first respiratory barrier against inhaled foreign matter and microorganisms. It is a lipoprotein complex comprising approximately 90 weight-% lipid of which phosphatidylcholine (PC) is the principal component. Importantly, the pulmonary surfactant is exceptionally rich in dipalmitoylphosphatidylcholine (DPPC) with a high main transition temperature (*T*_m_). Other major lipid components involve PCs with an unsaturated chain, phosphatidylglycerol (PG), and cholesterol.^3^ As to protein content, the pulmonary surfactant contains four specific surfactant proteins (SP-A, SP-B, SP-C, and SP-D) of which SP-A and SP-D are innate immune defence proteins, whereas SP-B and SP-C together with the phospholipids are crucial in sustaining the very low surface tension needed to avoid alveolar collapse, oedema, and lack of oxygenation. ^4^

Basic cellular and molecular biology research has suggested that early pulmonary surfactant dysfunction contributes to the high morbidity of coronaviruses.^2^ The ability to reduce the surface tension of pulmonary surfactant can be compromised due to decreased concentration of surfactant phospholipids and proteins, altered phospholipid composition, proteolysis and/or protein inhibition, as well as oxidative inactivation of lipids and proteins.^5–8^ Variations of these dysfunctional mechanisms have been reported in child and adult patients with ARDS. For instance, pulmonary surfactant metabolism studies in adult ARDS patients showed altered surfactant lipid composition: DPPC content was decreased, whereas the fractions of the surface tension-inactive unsaturated species were increased.^9^ Moreover, both the total amounts of PC and PG were decreased.^9^ Therefore, the possibility that early surfactant replacement therapy could be beneficial in preventing progression of disease severity is encouraging, given the established harmless profile of the pulmonary surfactant. Even though it is mainly indicated for prematurely born babies, surfactant replacement therapy has also been used in adult ARDS studies.^10–13^ This method,^14^ where exogenous surfactant is supplied into the lungs, is currently being tested in clinical trials to treat COVID-19 infected patients that require ventilator support.^15,16^ Accordingly, for the development of efficient surfactant replacements, we require a better understanding of the roles of the pulmonary surfactant components in lung mechanics.

Pulmonary surfactant forms a network of complex biological self-assemblying morphologies lining the alveoli. The distinctive structure formed by the pulmonary surfactant is a monolayer at the gas–alveolar epithelium liquid interface. In addition to lateral packing in the monolayer at the liquid–air interface, a fraction of the pulmonary surfactant is also likely folded from the interface into lipid bilayers or multilayers in the aqueous subphase,^17^ acting as lipid reservoirs.^18,19^ This monolayer is repeatedly compressed and expanded during breathing cycles; property that can only be withheld by a material with very peculiar viscoelastic properties.^20,21^ These reservoirs have also been suggested to participate in oxygen transfer from the inhaled air to circulation.^22^ It has been shown that the pulmonary surfactant monolayers and membranes exhibit phase behaviour that is believed to play roles in lung mechanics; tight packing of lipids with saturated lipid chains promotes its ability to lower surface tension, whereas lipids with unsaturated chains increase pulmonary surfactant fluidity and thus allow for its rapid adsorption to the air–water interface.^23–26^ Moreover, the phase behavior is also considered to regulate the functions of the surfactant proteins.^18,27^ However, the biophysical implications of the phase behavior are poorly understood, and a molecular view of the molecular organization would be extremely helpful in understanding the roles of different lipids and surfactant proteins in lung functionality enabled by pulmonary surfactant.

When compressed under the *T*_m_ of the corresponding membrane, single-component monolayers transition from a two-dimensional gas phase into a loosely packed and dynamic liquid-expanded (L_E_) phase. Further compression takes the monolayer to a brittle liquid-condensed (L_C_) phase, which eventually collapses, excreting matter into the aqueous subphase.^28–30^ The L_E_–L_C_ transition takes place through the formation of a coexistence phase characterized by a plateau in a surface pressure–area isotherm, which is manifested as observable small domains.^31–33^

The behavior of native pulmonary surfactant monolayers is quite different from such model systems. Still, fluorescence and Brewster angle microscopy reported visible domains in monolayers formed from lipid fractions of the pulmonary surfactant.^25,34^ These domains were suggested to be highly enriched in DPPC.^34^ Domains were also detected in vesicles created from the pulmonary surfactant, and also in the presence of the surfactant proteins.^24,25^ However, the surface pressure–area isotherms measured for pulmonary surfactant lipid fractions were not found to contain a plateau of any sort.^34^ Thus, the monolayer did not entirely transition to the L_C_ phase at any pressure, indicating that no L_E_–L_C_ coexistence was present either. Presumably, the observed domains did not present an equilibrium phase. Indeed, the composition of the pulmonary surfactant seems to be adjusted so that the mixture is barely fluid at body temperature,^24,35^ which can lead to distinct behavior of its lipid components. By doping the pulmonary surfactant lipid fraction with additional DPPC, the *T*_m_ can be gradually increased, and a coexistence plateau becomes eventually visible.^34^ These observations suggest that the behavior of the pulmonary surfactant may be characterized by transient heterogeneity arising from critical fluctuations,^36^ where the behaviour of each lipid type is determined by its *T*_m_ value.

These research topics are difficult to study at the molecular level due to the limited resolution of available experimental techniques. Molecular simulations are often helpful in resolving such aspects down to the atomistic level.^37^ However, previous simulation studies on multi-component monolayers are relatively scarce.^17,29,38–42^ While previous simulations have generated a lot of insight to better understand the behavior of surfactant monolayers, they have been haunted by the insufficiently accurate description of the physical behavior of the lipids at the interface, which is largely due to water models whose quality in describing phenomena at water-monolayer–air interfacial regions is not sufficiently high,^33,39,43,44^ and due to incorrect description of the driving forces behind the formation of heterogeneous membranes.^45^

To overcome these limitations and provide a detailed and accurate picture of the molecular-level organization of surfactant monolayers, we used extensive and state-of-the-art all-atom molecular dynamics (MD) simulations of model systems that were validated with experimental Langmuir trough data. The composition of our model monolayers was chosen to match the composition of the protein-free pulmonary surfactant.^25^ These model systems were simulated at various compression states and temperatures, matching the conditions studied in experiments. By using our recently developed protocol for performing simulations of lipid monolayers at the air–water interface,^33,46^ we were able to reach quantitative agreement with experimental surface pressure–area isotherms.

The simulations predicted the presence of lateral heterogeneity characterized by domain formation, which we confirmed by atomic force microscopy (AFM) imaging. Upon compression, we found the appearance of transient L_C_-like domains. Upon further compression these domains were found to aggregate to form large ordered regions. Thanks to the atomistic detail available in the simulations, we were able to draw conclusions as to the physical and chemical properties of the nanoscale domains. Interestingly, we observed that the domains are not substantially enriched by any lipid type, indicating that the peculiar viscoelastic properties of the pulmonary surfactant arise from the collective behavior of the mixture rather than from the features of its independent lipid components. Our findings help understand lung mechanics and thus guide the development of strategies to tackle lung conditions, while also paving the methodological way for future studies of the pulmonary surfactant.

## Methods

### Atomistic Molecular Dynamics Simulations

Until recently, the poor description of surface tension by the commonly used water models^43,47^ has prevented quantitative simulation studies of lipid monolayers at the air–water interface. However, very recently we demonstrated that a combination of the four-point OPC water model^48^ and the CHARMM36 lipid model^49^ reproduces experimental surface pressure– area isotherms in single-component 1-palmitoyl-2-oleoylphosphatidylcholine (POPC) and DPPC monolayers,^33^ and the agreement is largely quantitative. In this study, we extend this approach to quaternary lipid monolayers whose composition was chosen to match the composition of the protein-free pulmonary surfactant^3^ as accurately as possible. In practice, our systems contained 60 mol% DPPC, 20 mol% POPC, 10 mol% 1-palmitoyl-2-oleoylphosphatidylglycerol (POPG), and 10 mol% cholesterol. The systems were set up as follows. Two monolayers, separated by a water slab and each containing 169 lipids were first set up at an area per lipid (APL) equal to 50 Å^2^. Next, these monolayers were either expanded or compressed during a 10 ns simulation to an average APL value of 100 or 40 Å^2^, respectively, using the `MOVINGRESTRAINT` and `CELL` keywords in the PLUMED 2.2 package.^50^ Structures at a total of 17 APL values were extracted and used as initial structures for the production monolayer simulations in the NVT ensemble for 1 µs each. All simulations were performed at both 298 and 310 K, and additional repeats as well as larger simulations of up to 3042 lipids were performed to provide further validation for the results.

The CHARMM36 model for phospholipids^49^ and cholesterol^51^ was used together with the four-point OPC water model.^48^ The simulations were performed with the version 5.1.x of the GROMACS simulation package,^52^ and the recommended simulation parameters for the CHARMM36 force field^53^ were used to reproduce realistic monolayer behavior.^33^ Details on the simulation setups and on the simulation methodology are provided in the Supporting Information (SI).

### Analyses of MD Simulations

#### Surface Pressure–Area Isotherms

Monolayer behavior was characterized and compared to experiments using surface pressure– area isotherms. Monolayer surface pressure Π at an APL of *A* was calculated as Π (*A*) = γ_0_ − γ(*A*), where γ and γ_0_ are the surface tensions of the monolayer-covered and plain air– water interfaces, respectively. Here, the surface tensions are obtained from the pressure components along the monolayer plane (*P*_L_ = (*P*_*xx*_ + *P*_*yy*_) */*2) and normal to it (*P*_N_) as γ = *L*_*z*_ × (*P*_N_ − *P*_L_) */*2, where *L*_*z*_ is the simulation box size normal to the monolayer plane, and a factor of two indicates the presence of two water–air interfaces in the simulation box. The values were obtained with the `gmx energy` tool provided with GROMACS.^52^

#### Detection of Monolayer Domains

Domains packed like in the L_C_ phase were detected by clustering lipid chains and cholesterols based on their packing in the monolayer plane. The 10^th^ carbons in the lipid chains and the C14 atom in cholesterol ring were included in the clustering that used the DBSCAN algorithm.^54^ The chosen atoms in the lipid chains capture well the hexagonal packing in ordered structures,^55^ and the C14 carbon of cholesterol resides at the same depth. This carbon is part of both the five-member (D) and six-member (C) rings. For DBSCAN, we used a cut-off of 0.71 nm and a minimum neighbour count of 6. The cut-off was set to the distance at which the first minimum appears in the radial distribution function of the clustered atoms. The clusters were considered to be part of the L_C_-like domain. This clustering was performed on conformations separated by 1 ns. The L_C_-like fraction, number of individual clusters, and the largest cluster size were extracted for each conformation. All these quantities were then averaged over the trajectory, and their standard deviations were used as error estimates. Pattern-matching was used to find all residence events in the L_C_ clusters. These times were histogrammed and fitted with a power law with an exponent of *b*.

### Contact Fraction

Lateral demixing of lipids with unsaturated and saturated chains was quantified by the contact fraction. Following Ref. 56, we defined the contact fraction as *f*_mix_ = *c*_US−S_*/*(*c*_US−US_ + *c*_US−S_), where *c*_US−S_ is the number of contacts between the lipids with an unsaturated chain (POPC and POPG) and the lipids with saturated chains (DPPC). For a contact, the phosphorus atoms of the lipids had to be within 1.1 nm from each other. We extracted contact data using the `gmx mindist` tool every 1 ns and calculated the average values over the last 500 ns of the simulations for both monolayers. We plot the mean of these two average values, whereas their difference serves as an error estimate.

#### Cholesterol Clusters

Cholesterols were considered to be part of a cholesterol cluster if they were in contact with at least one other cholesterol molecule. We used a cut-off of 0.94 nm based on the first minimum in the cholesterol–cholesterol radial distribution function. All residence events were used in the calculation of the probability distribution of cholesterol cluster sizes. The distances were measured from the centers of mass of the cholesterol molecules.

#### Diffusion Coefficients

Diffusion coefficients were calculated to characterize monolayer dynamics and to detect changes in monolayer packing. The diffusion coefficients were extracted from center-of-mass (COM) trajectories. The motion of lipids with respect to the movement of the monolayer as a whole was analyzed to eliminate possible artifacts due to monolayer drift. The diffusion coefficients were extracted from linear fits to time- and ensemble-averaged mean-squared displacement at lag times between 10 and 100 ns. Two values were extracted — one from each monolayer — and averaged, and their difference served as an error estimate. The GROMACS tool `gmx msd` was used, and the COM trajectories were generated using `gmx traj`.

#### Monolayer Thickness

Monolayer thickness was used to couple AFM height profiles of heterogeneous membranes to the lateral packing within the domains. This thickness was estimated from density profiles calculated for each lipid type along the monolayer normal (*z*) using the GROMACS tool `gmx density`. These profiles were aligned at the phosphorus peak of DPPC, and the lipid was set to begin and end at *z* values where its density crossed 5% of its maximum value.

#### Lipid Chain Tilt

Lipid chain tilt was used to characterize persistent L_C_-like packing in the monolayers. The tilt angle of lipid chains was calculated as the angle between the *z* axis and the vector joining the 1^st^ and 16^th^ carbons in the fatty acid chains of phospholipids using the GROMACS tool `gmx gangle`. The angle distributions were averaged over both chains and both monolayers, and the distributions were fitted with a Gaussian. The location of its maximum and the variance were used as the mean tilt angle and its error estimate, respectively.

### Langmuir Trough Measurements

#### Langmuir–Wilhelmy Compression Isotherms

By means of a specially designed ribbon Langmuir–Wilhelmy trough (NIMA Technology, UK), compression isotherm assays were performed and surface pressures as a function of molecular area were obtained at constant temperature, as described previously.^57^ The lipid mixture was identical to that employed in the simulations (see above). The employed Langmuir–Wilhelmy trough has a maximum area of 312 cm^2^ and a minimum of 54 cm^2^ and instead of the canonical rigid barriers uses a continuous Teflon-coated ribbon that by moving symmetrically reduces the available area in the aqueous surface, and therefore compresses the confined sample to carry out the compression isotherm. The pressure is recorded using a electronical pressure sensor and a piece of cellulose and employing the Wilhelmy plate technique, with an estimated error of ±1 mN/m among different isotherms, that were assessed at least in triplicate at 298 K. The compression isotherm displayed in Fig. 1 shows the first compression of 44.1 nmol of a DPPC/POPC/POPG/Chol mixture of mimetic surfactant monolayer, spread on an aqueous-buffered subphase composed of 5 mM Tris, pH 7.4, 150 mM NaCl. After placing the lipid mixture onto the aqueous subphase surface, the generated lipid layer was left to equilibrate for 10 min, so the solvents could evaporate, and the lipids organise and disperse at the interface. Thereafter, the narrowest area was achieved by compressing the monolayer at a constant speed of 150 cm^2^/min. This compression speed was set to reflect the functional quality of the assessed surfactant, by recapitulating as much as possible the dynamic context of native pulmonary surfactant in the human organism, and allow reaching the highest surface pressure possible.

**Figure 1:**
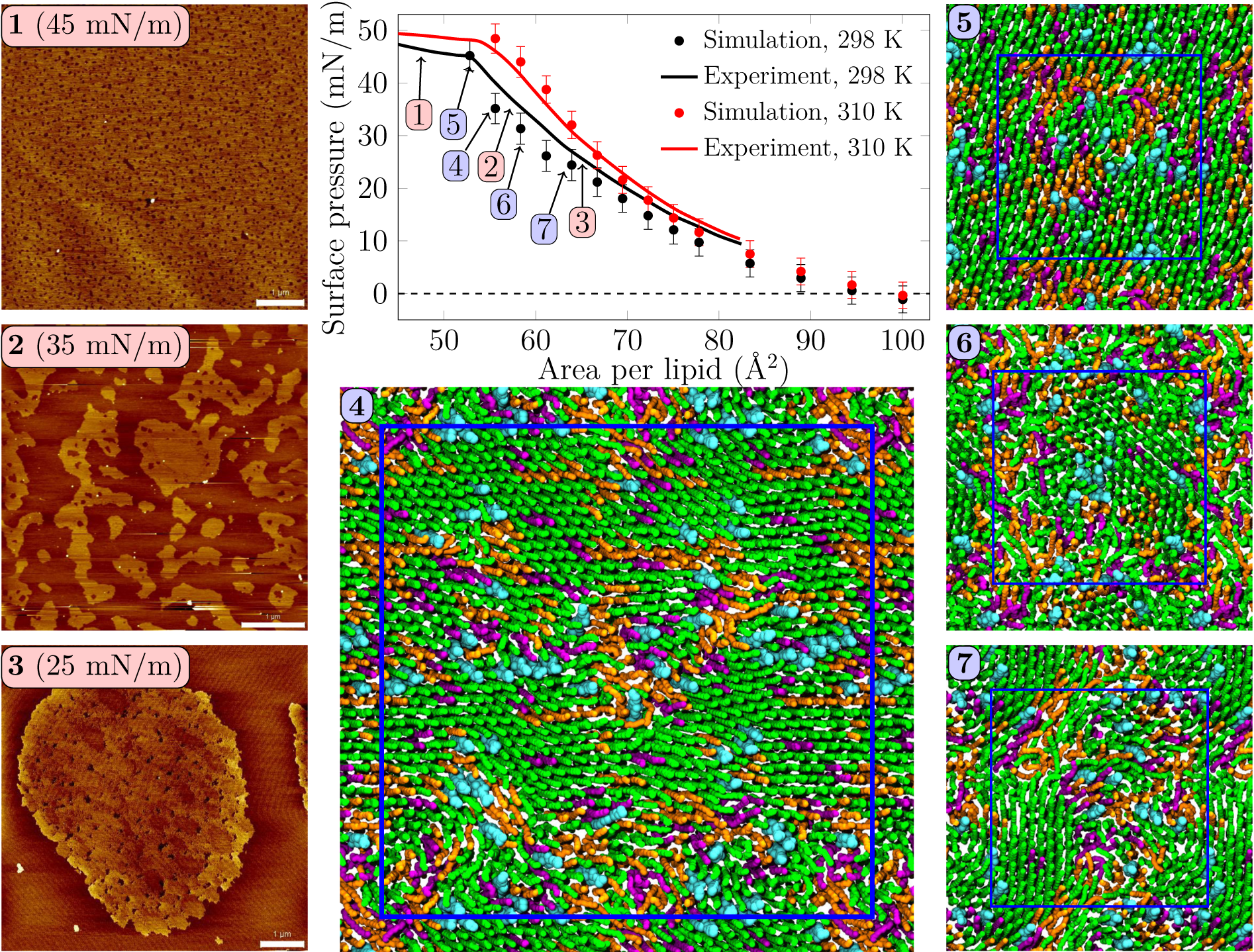
Surface pressure–area isotherms from simulations and experiments, with AFM images and simulation snapshots taken from selected surface pressures. Scale bar 1 µm. At 25 mN/m, (**3**) one can identify small irregularly shaped domains with visible nanostructures inside the domains. At 35 mN/m, (**2**) the domains start to fuse and lose their rounded shapes. In the collapse plateau at 45 mN/m, (**1**) the domains are no longer visible in the AFM images, and some lipids are squeezed out from the monolayer film. Clearly visible transient or persistent heterogeneity can also be observed in the simulation snapshots (**4– 7**) of monolayers at different surface pressures. Here, **4** was extracted from a 4-fold larger monolayer than **5–7**. Only the data points below the experimental collapse pressure are shown in the isotherm, whereas all simulated points with all of the independent repetitions are shown in Fig. S1.

#### Surface Dilatational Rheology

We again used a lipid composition identical to the one in simulations (see above). All these lipids were obtained from Avanti Polar Lipids (Alabaster, AL). Lipids were spread in 2 mM chloroform solution on the surface of a deionized water subphase (Milli-Q, Millipore, Bedford, MA) in a Langmuir trough (KSV Minitrough, Espoo, Finland) with a platinum Wilhelmy plate. The dilatational rheology of the monolayer was studied using the oscillating barrier method. The film was first compressed to the desired surface pressure, after which sinusoidal area compressions with an amplitude of 1% were performed at a frequency of 10, 50, 100, or 200 mHz and the changes in surface pressure in response to the oscillations were recorded to determine the surface dilatational modulus. Each measurement was repeated two times, and an average dilatational modulus was calculated over frequency, since the dilatational modulus was found to be constant with respect to frequency. All measurements were conducted at room temperature (297 K).

### Atomic Force Microscopy Imaging

#### Langmuir–Blodgett Transfered Monolayers

Lipid monolayers were transferred onto mica as described in Ref. 58. For the monolayer preparation, the sample was spread onto the air–water interface until the minimum surface pressure of ∼0–1 mN/m was observed. After 10 min of monolayer equilibration, the film was compressed until the desired surface pressure was reached (from 15 to 45 mN/m, in 5 mN/m steps), at a compression speed of 50 cm^2^/min. Before the transfer was started, the film was again equilibrated for 5 min at constant pressure. The monolayers were finally deposited in a freshly cleaved muscovite mica substrate (Plano GmbH, Wetzlar, Germany) that had been previously submerged. The lifting device—to which the mica substrate was fixed–was raised in the vertical plane out of the buffered aqueous subphase at a speed of 10 mm/min at constant pressure. In all experiment modalities three independent experiments were carried out, as a minimum, and up to 10 images were taken and analysed.

#### Atomic Force Microscopy

Langmuir–Blodgett supported monolayers’ topographical images were taken using an atomic force microscope (JPK NanoWizard, JPK Instruments, Berlin, Germany), employing in both cases Silicon-SPM cantilevers (Nanosensors, NanoWorld AG, Neuchatel, Switzerland). The AC mode in air was selected for monolayers. The scan rate was ∼1 Hz for all AFM images. At least 3 different supported mono- and bilayer systems were assessed, and each sample was imaged on a minimum of three different positions. Image processing of AFM data was done using the SPIP software package as in Ref. 25 (Image Metrology, Hørsholm, Denmark).

## Results

### Monolayers Display Lateral Heterogeneity Without a Coexistence Plateau

Surface pressure–area isotherm is a key quantity representing monolayer behavior at the water–air interface, and it is readily extracted from both Langmuir trough measurements and computer simulations. These isotherms for the quaternary pulmonary surfactant lipid monolayers at 298 and 310 K are shown in Figure 1.

The isotherms from experiments and simulations are in nearly quantitative agreement. Although no plateau indicating L_C_/L_E_ coexistence was visible in the experimental isotherm, our AFM measurement displayed heterogeneity in lipid packing. The AFM images taken at 298 K (Fig. 1, panels **1–3**, and Fig. S2–S5) revealed surface pressure-dependent formation of a heterogeneous monolayer with thinner and thicker regions. Starting from low surface pressure (higher APL), at 25 mN/m (Fig. 1, image **3**) the observed domains were rounded yet irregular in shape. The insides of these domains are heterogeneous with visible nanostructures, indicated also by our height profile studies of the domains (see Fig. 2 and Figs. S6–S9). Similar heterogeneity was also evident in the simulations (Fig. 1, panels **4–7**), where domains were readily seen at surface pressures above 25 mN/m, or at APL below 65 Å^2^ at 298 K. No L_C_/L_E_ coexistence plateau was observed in the simulated isotherms either (Fig. 1 and Fig. S1), which was further verified by running independent replica simulations at selected APLs. Furthermore, by running independent repetitions at selected areas and larger monolayer systems, we also verified that the initial monolayer configuration or the finite-size effects did not affect the results (Fig. S1). The movies at DOI:10.6084/m9.figshare.12612317 as well as the additional snapshots of the simulated monolayers at 298 K and 310 K given in Figs. S10 and S11, respectively, clearly demonstrate heterogeneity in lateral organization. Based on our simulations at 298 K (Fig. 2C), the monolayers at APLs equal to 50 (corresponding to the very packed state with major regions in an L_C_-like arrangement) and 90 Å^2^(corresponding to the L_E_ phase) extend 2.5 and 1.8 nm towards the air phase, respectively, calculated from the plane defined by the DPPC phosphorus atoms (see Methods). This difference of 0.7 nm is exactly the difference observed in the height profile study in the AFM experiments (Fig. 2A and B), clearly indicating that the measured thicker and thinner regions in the monolayers correspond to L_E_- and L_C_-like regions, respectively.

**Figure 2:**
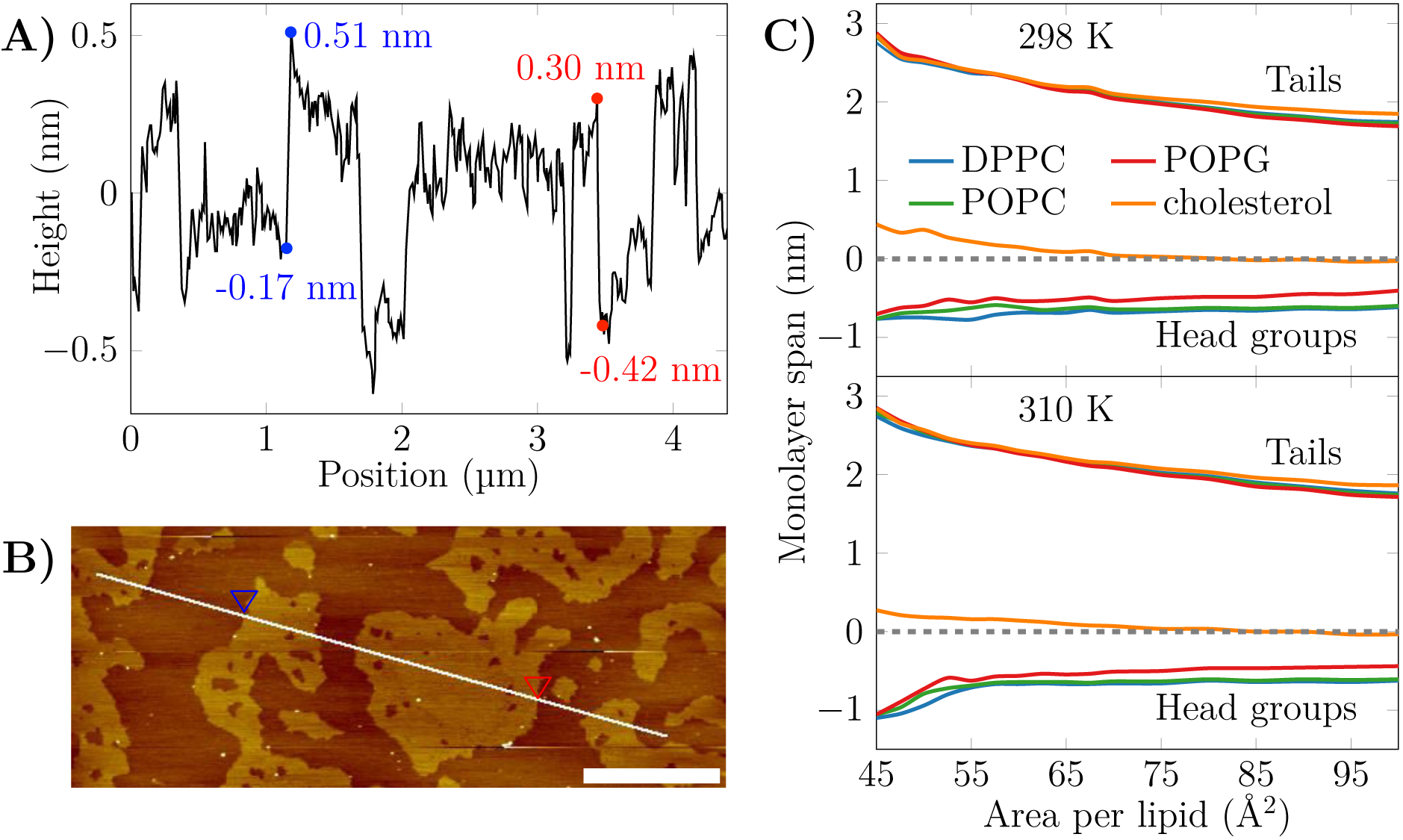
The AFM height profile at 35 mN/m and the average monolayer thickness calculated from the simulations. A) The AFM height profile results show a difference of approximately 0.7 nm at the border of the two phases at the selected positions. B) The path of the measurement. Two points marked with the blue and red triangles indicate selected positions at the border of the L_C_- and L_E_-like regions. The scale bar at the lower right corner equals to 1 µm. C) The average thickness of the monolayer in the simulations. The dashed line indicates the position of maximum of phosphorus density of DPPC. The lines indicate the span of lipids from tails to head groups. At each extreme, the line shows the location where the lipid density drops below 5% of its maximum value.

The extent of the L_C_-like region grows as the surface pressure of the monolayer is increased, as would also be expected during a proper phase coexistence region,^33^ although the surface pressure does not remain constant here based on the isotherms shown in Fig. 1. From AFM figures, at 35 mN/m (Fig. 1, image **2**) the L_C_ domains seem to fuse together to form more irregular shapes, but the overall heterogeneity is clearly visible. Here, the coalescence of the domains was likely also limited by the slow diffusion within the compressed monolayers. Indeed, our simulations predict that compression from a state with APL equal to 80 Å^2^ to the one with APL equal to 50 Å^2^ slows lipid diffusion by 1–2 orders of magnitude (Fig. 3A). The nanostructures within the domains correspond to the L_C_-like ones measured at 25 mN/m (see SI). At 45 mN/m (Fig. 1, image **1** below APL 55 Å^2^) and 55 mN/m (Fig. S5 in the SI) we have already reached the monolayer equilibrium collapse plateau seen in the experimental surface pressure–area isotherms.^31^ It should be noted that the monolayers in the alveoli are expected to reach surface tension values close to 0 mN/m (corresponding to surface pressure of ∼70 mN/m). However, in exeriments this is only achieved by very rapid compression,^59^ or by the use of bubble surfactometer methods.^60^ In the collapse region, the domains are completely merged with a continuous network of small holes, forming a sponge-like collapsed structure at the boundary of the air–liquid interface. Further on, the protruding regions seen in the AFM images at surface pressures above the collapse plateau are collapsed material, excluded from the interfacial film into the water phase. This is verified by the height profiles (Fig. S9 in the SI), which show height differences of up to 10 nm, which is clearly above the difference observed in the simulations between monolayers at different APLs (Fig. 2C). Increasing the temperature to 310 K in the simulations renders the monolayer more fluid and the heterogeneity is visible at surface pressures above 30 mN/m, or APL below 60 Å^2^ (Fig. S11 in the SI).

**Figure 3:**
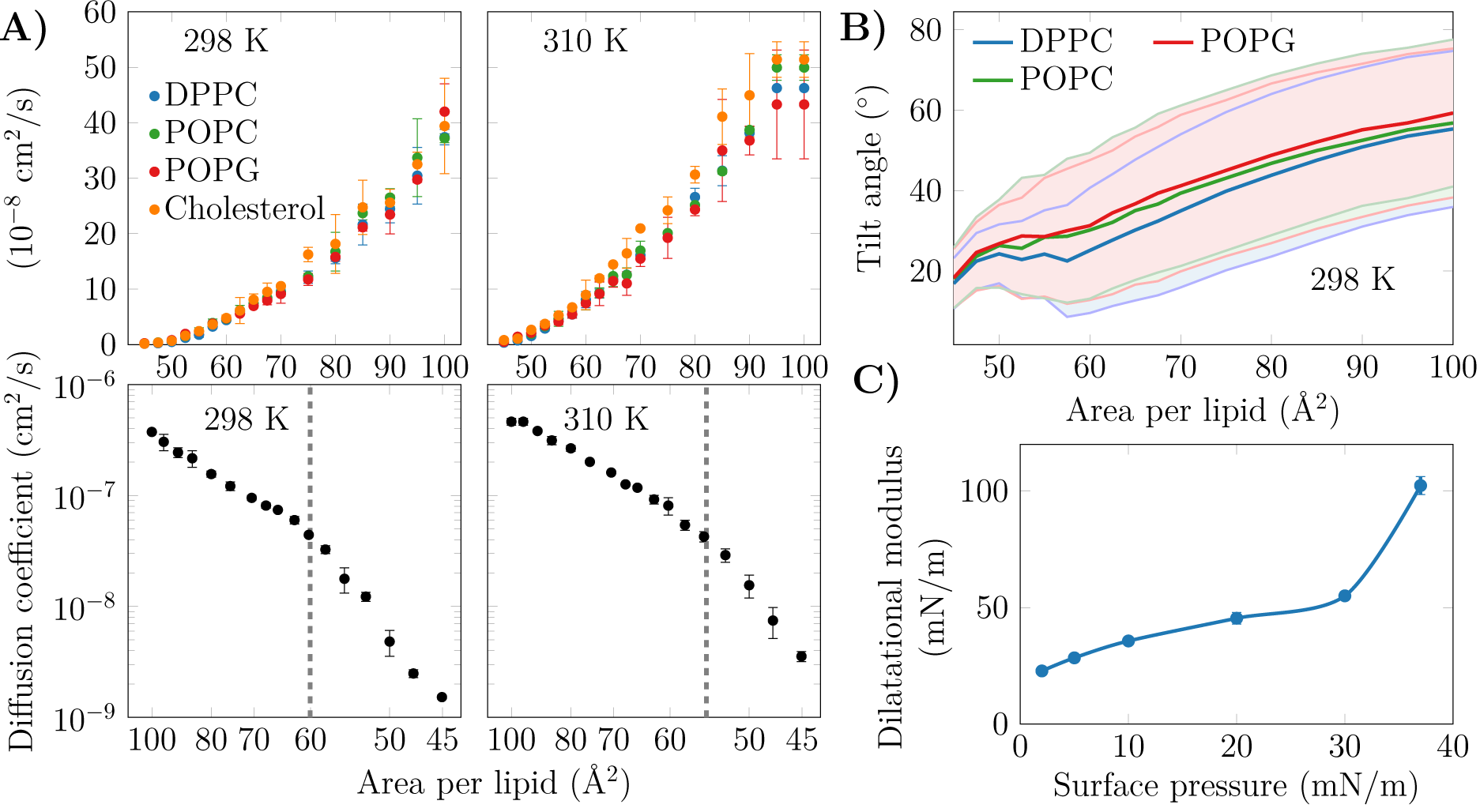
A) Diffusion coefficients *D* of the lipids in the monolayers. Top row: The diffusion coefficients, shown in linear scale as a function of APL, show no signs of a phase transition at either studied temperature. Bottom row: *D*, in logarithmic scale, shown as a function of 1/APL displays two slopes with a crossover at APL≈60 Å^2^ (298 K) or ≈55 Å^2^ (310 K), highlighted with a dashed gray line. B) Tilt angles of the phopsholipid chains in the mono-layers at 298 K. The shaded areas, bordered by dim lines, show the standard deviation. Data for 310 K is provided in Fig. S12 in the SI. C) Dilatational modulus of the monolayer as a function of surface pressure at 297 K.

Concluding, both AFM measurements and computer simulations revealed lateral heterogeneity in a large range of APL values, despite the lack of a plateau in the surface pressure–area isotherm. Together, simulations and AFM imaging suggest that the structure of these domains resembles the L_C_ phase.

### Condensed Nanodomains Affect the Structural and Dynamic Properties of the Monolayer

As no coexistence plateau is observed in the isotherms obtained from experiments and simulations, it is worth asking whether the heterogeneity observed by AFM and simulations manifests itself in some other monolayer properties. To this end, we measured the dilatational moduli of the monolayer at various surface pressures using the oscillating barrier approach. As shown in Fig. 3C, we observe little change in this modulus up to a surface pressure of 30 mN/m. This signals that the monolayer area can be changed relatively easily, possibly due to a rapid reorganization of lipids in the membrane plane. However, after 30 mN/m — corresponding roughly to APL equal to 60 Å^2^ based on the isotherms in Fig. 1 — the modulus grows rapidly. This indicates that there are structural changes in the monolayer, perhaps due to the formation of continuous ordered regions.

On the simulation side, we analyzed the diffusion of lipids at different APLs. The diffusion coefficients, *D*, are shown in the top row of Fig. 3A. The curves demonstrate what seems to be a continuous and smooth increase in diffusion coefficient upon an increase in APL. However, as shown in the bottom row of Fig. 3A, a plot of log(*D*) versus 1/APL does not reveal a single slope, as expected for monolayers without a plateau in the pressure–area isotherm, such as DLPC at 294 K.^61,62^ Instead, we observed two distinct slopes with crossovers at 60 and 55 Å^2^ for DPPC at 298 and 310 K, respectively. The presentation of the diffusion data used herein is based on the obvious idea that the diffusion of lipids depends on their degree of packing. This presentation resembles that used to interpret diffusion in terms of the free volume theory.^63^ However, due to various issues in applying this theory to diffusion in lipid monolayers,^40^ we refrain from interpreting the diffusion data further within this framework. Yet, we acknowledge a clear change in the trend of the APL-dependence of diffusion coefficients.

Let us move on to a structural property extracted from the simulations. The tilt angle distributions at 310 K, shown in Fig. S12 in the SI, show a continuous increase in the chain tilt upon increasing area. However, as show in Fig. 3B, at 298 K and between APLs of 45 and 60 Å^2^, the chains remain tilted at an angle of ∼25^°^. A similar tilt angle of ∼25^°^ was also reported for DPPC monolayers in the L_C_ phase using both X-ray diffraction^64^ and vibrational sum frequency generation spectroscopy.^65^ This indicates that at 298 K, the monolayer retains regions of L_C_-like packing with a characteristic chain tilt until relatively

large APL values, and this structural feature is coupled to a diffusion mode that differs from that observed at larger APL values. It is worth noting that the long-range orientational order of the tilted L_C_-like domains^66^ is also clearly evident in Fig. 1.

Concluding, in both experiments and simulations we systematically observe a a sudden change in monolayer properties at a certain compression level, despite the isotherms showing no coexistence plateau.

### Condensed Domains Display Little Enrichment in DPPC and Cholesterol

With the presence of L_C_-like domains evident in both simulations and experiments, we next use the simulation data to study the composition of the domains. The leftmost panels in Fig. 4A show the fraction of phospholipid chains and cholesterols in L_C_-like domains obtained from clustering analyses (see Methods). The data are resolved by lipid type and shown in a cumulative manner so that the total L_C_-like fraction can be read from the plot. At low APL values, almost the entire monolayer is part of the L_C_-like region, whereas at large APL only transient clusters of a few lipids are observed. Still, more than half of the phospholipid chains and cholesterols are part of an L_C_-like domain up to APL values of ∼65 Å^2^. Interestingly, temperature has little effect on the calculated L_C_ fractions, since very similar domain compositions are extracted from the simulations at room and body temperatures.

**Figure 4:**
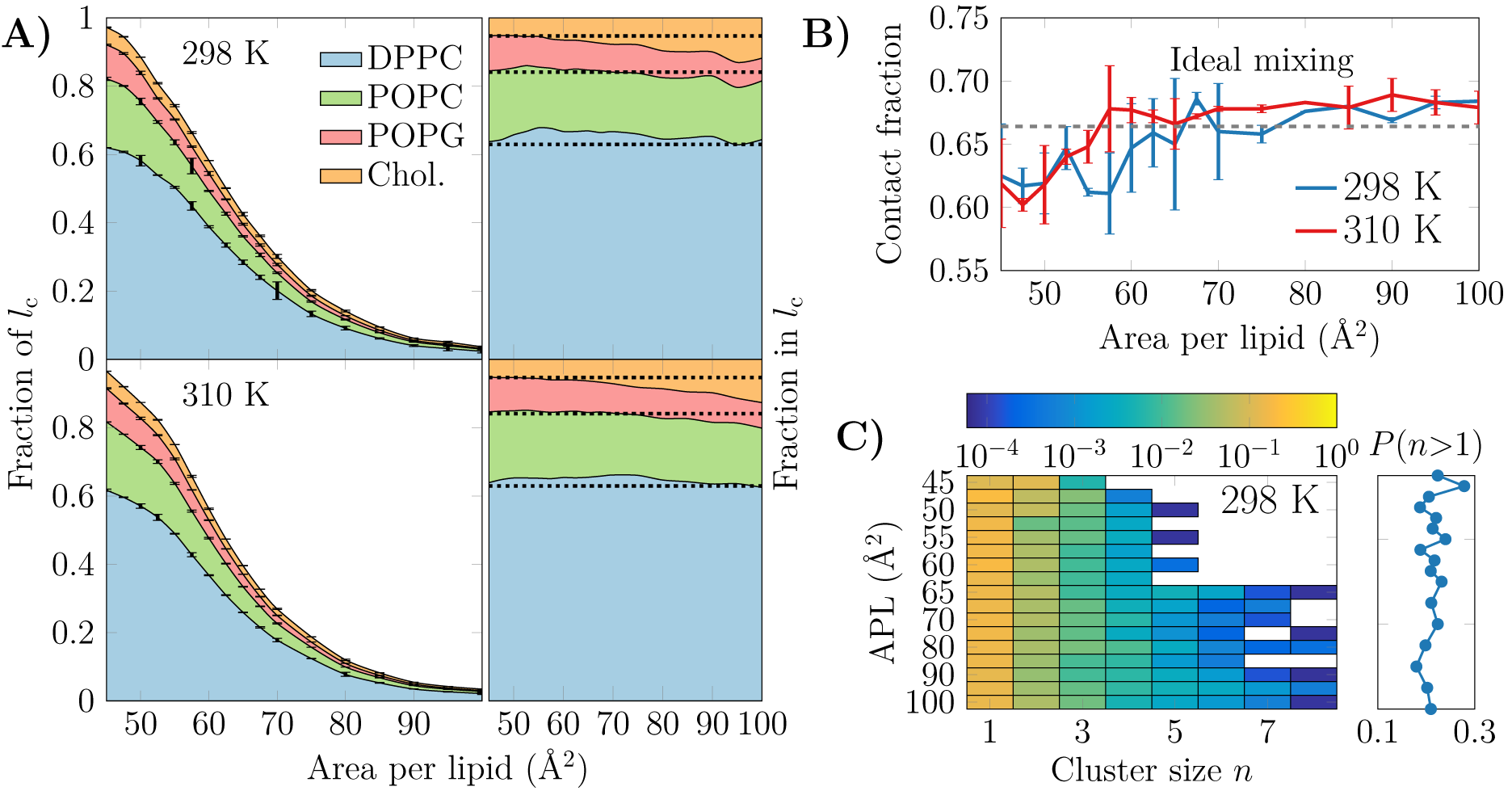
A) The extent and composition of the L_C_-like regions. On the left, the fraction of phospholipid chains and cholesterols in the L_C_-like regions is shown for all lipid types. These curves sum to the total fraction of the L_C_-like region. The error bars show the differences calculated for the two monolayers in the simulation system. On the right, the fraction of lipids that make up the L_C_-like regions at different APLs, *i*.*e*. in this plot the curves always sum to 1. The dashed lines show the values expected based on the overall monolayer lipid composition. B) Mean contact fraction during the last 500 ns between DPPC and the lipids with unsaturated chains (POPC and POPG). Ideal mixing is highlighted by a dashed line at 0.66, whereas values smaller than that indicate a preference to demix. C) Left: 2-dimensional probability distributions of cholesterol molecules residing in a cluster with at least one other cholesterol at all simulated APLs at 298 K. Note that the color bar is in logarithmic scale. Right: The sum of probabilities of *n>* 1, *i*.*e*. the probability for cholesterol to be in a cluster with size larger than one molecule. Data for simulations at 310 K are given in Fig. S13 in the SI.

It is clear from Fig. 4A that DPPC with its two saturated chains makes up the majority of the L_C_-like regions, regardless of the surface pressure and temperature. This is natural, since DPPC is also the most prevalent lipid type in our simulations. For a more detailed look into the composition of the L_C_-like domains, we plot the fraction of lipids in these regions in the right panels of Fig. 4A. Here, the dashed lines highlight the overall fractions of the different components in the system. At very low APL, essentially the entire monolayer is detected by the clustering algorithm to be part of a L_C_-like region, and therefore the fractions of lipids in this region agree with their fractions in the system. However, at intermediate APLs between 50 and 80 Å^2^, DPPC is present in the L_C_-like domain slightly more than expected by its fraction in the system. The fraction of POPC within the L_C_-like regions is fairly constant over all APLs, whereas those of POPG and cholesterol change more noticeably while moving towards larger APLs; the fraction of POPG decreases while that of cholesterol increases in the shrinking and more transient L_C_-like regions. Indeed, at large APLs, cholesterol seems to be a key player in the formation of condensed lipid clusters.

It is quite surprising that the L_C_-like domains have so little preference for any lipid type. We therefore also analyzed whether the lipids underwent lateral demixing during the simulation. The mean contact fractions between lipids with unsaturated and saturated chains in all simulations are shown in Fig. 4B. The dashed line at 0.66 represents ideal mixing of DPPC with the lipids with unsaturated chains (POPC and POPG). It is evident from Fig. 4B that there is generally little preference for lipid demixing, yet at smaller APL values there is a small tendency for DPPC lipids to reside with other DPPC molecules. The APL values below which lipid mixing deviates from ideal behavior are again close to 55 and 60 Å^2^, where rapid changes in the APL-dependence of many quantities were observed in the last section. Curiously, temperature has again very little effect on this demixing. Finally, it is worth pointing out that we also studied the time evolution of the contact fractions of the systems at different APL values but found no systematic tendency for any of the monolayers to demix during the simulation time.

We also studied whether cholesterol has a tendency to cluster in the monolayers. The probability of a cholesterol to be in a cluster of a given size is shown in the left panel in Fig. 4C for the simulations at 298 K, whereas the data for simulations at 310 K are given in Fig. S13 in the SI. The right panel shows the sum of probabilities with *n* ≥ 2, corresponding to the probability that a cholesterol molecule is part of a cluster with at least one other cholesterol molecule.

It is evident from Fig. 4C that all distributions peak at 1, indicating that most cholesterol molecules are on average not in contact with other cholesterol molecules. Still, the probability of a cholesterol to be in a cluster with size larger than one is ∼20% in all monolayers (right panel in Fig. 4C). Curiously, at APLs 65 Å^2^ and larger, clusters of more than 5 cholesterol molecules sometimes appear. Still, the probability of cholesterol clusters larger than one shows no clear dependence on APL, although not surprisingly, the chance of finding cholesterol clusters is highest in the monolayers with the smallest APLs.

To analyze the distribution of the domains in the monolayer, we show in Fig. 5A the number of condensed clusters as well as the fraction of the system occupied by the largest cluster. Between APLs of 45 and 60 Å^2^ the size of the largest cluster drops steeply from a coverage of ∼80% of the lipids to a mere ∼10%. At the same time, the number of L_C_ clusters increases from a few to 15–20, and these maxima are reached at an APL equal to 60 Å^2^ (298 K) or slightly smaller (310 K). These APL values agree well with those at which changes in trends as a function of APL were observed in many properties in the previous section. The melting of the largest cluster is coupled to the formation of many smaller clusters. Upon an increase in APL beyond 60 Å^2^, the number of clusters begins to decrease again, until only a few are detected in the pure L_E_ phase. Moreover, these few are very small, since the largest one occupies only 2% of the lipids, or less, indicating that it is likely a false positive due to density fluctuations and the used clustering algorithm. It is also noteworthy that at 298 K and between 45 and 70 Å^2^, there are slightly fewer domains as compared to the case at 310 K. However, the largest domain at 298 K is respectively larger in this interval of APLs.

**Figure 5:**
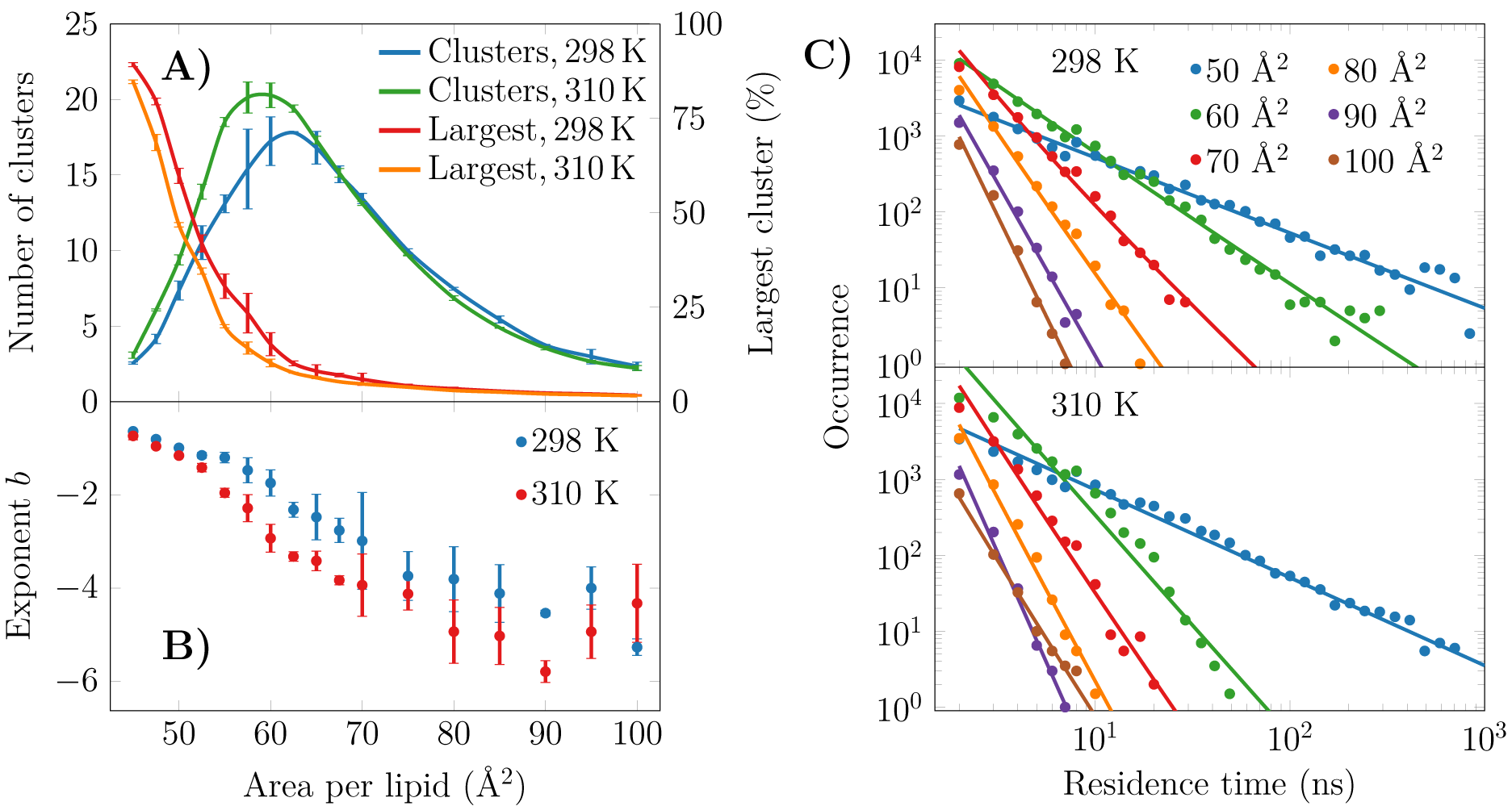
Spatial and temporal extent of the L_C_-like regions. A) The numbers of clusters in monolayers at 298 and 310 K are shown in green and blue, respectively, whereas the largest clusters in these systems are indicated red and orange, respectively. B) The exponents *b* of the fits in the form of *P* (*τ*)= *a* × *τ* ^*b*^ to the residence time distributions in panel C). The error bars show the differences calculated for the two monolayers in the simulation system. C) The lifetime distributions of the L_C_-like domains for both 298 K (top) and 310 K (bottom). The solid lines show the power law fits.

To evaluate whether the observed domains are transient or long-lived, we analyzed the distributions of residence time of the lipid chains in the L_C_-like regions. The residence time distributions for selected monolayers are shown in Fig. 5C. These distributions clearly show power law scaling, *i*.*e*. the probability *P* of a binding time *τ* follows *P* (*τ*) = *a* × *τ* ^*b*^. In the log–log scale, we extracted exponents *b* from a linear fit, and these exponents are shown in Fig. 5B. The larger in absolute value of *b*, the faster the distribution decays, *i*.*e*. the less lipids reside for extended times in the L_C_-like clusters. The exponents decrease upon increasing APL indicating that the residence times of lipids in the L_C_-like regions decrease. At an APL of 50 Å^2^, there are more than 100 events where a lipid stays in the L_C_-like domain for more than 100 ns, and some lipids stay there throughout the entire simulation. Interestingly, the exponents start to rapidly decay at APLs of 60 Å^2^ (298 K) or 55 Å^2^ (310 K), indicating that long-lived domains are replaced by more dynamic heterogeneity. These APL values, again, agree well with those at which significant changes are observed in other monolayer properties (see previous section). At APLs smaller than 55 Å^2^, the distributions at the two temperatures are very similar due to the large extent of the L_C_-like regions, whereas there are clear differences in the APL range between 55 and 75 Å^2^. Namely, the residence times for monolayers at 298 K are drastically longer. At areas above 75 Å^2^, the L_C_-like clusters are extremely dynamic and the longest residence times do not reach much over 10 ns regardless of the temperature.

## Discussion

We first extracted surface pressure–area isotherms from simulations and experiments. As these isotherms did not display a coexistence plateau, we performed AFM experiments that demonstrated the presence of sub-micrometer sized domains (Figs. 1 and S2–S5). The height profiles (Fig. 2B and Figs. S6–S9) and the thickness profiles obtained from our simulations (Fig. 2C) indicated that the domains resemble the L_C_ phase, whereas the remainder of the monolayer is in the L_E_ phase. A detailed investigation of the atomistic simulation data on the condensed domains showed that at large APL (*>*60 Å^2^) these domains consisted of transient, isolated islands. As long as the condensed domains are “cushioned” by the surrounding L_E_ phase, the macroscopic properties of the monolayer such as compressibility resemble those of the L_E_ phase. Interestingly, a similar observation emerged from a recent coarse-grained simulation study on the elastic properties of lipid bilayers;^67^ it was shown indeed that, in the presence of phase separation, bilayer elastic properties (including both the bending modulus and the area compressibility modulus) are close to the values calculated for the softer component. The present investigation extends the previous conclusion to monolayers not showing proper phase separation, but still presenting structural heterogeneity in the form of dynamic nano-sized domains. At smaller APL (*<*60 Å^2^) the condensed domains become stable and form a continuous meshwork, which explains the sudden increase in rigidity of the monolayer indicated by the change in the dilatational modulus in Langmuir trough measurements (Fig. 3C). The shift from transient to stable condensed domains was also accompanied by a change in the lateral diffusion dynamics in the monolayer, as well as in the lipid mixing properties (Fig. 4B) and in the lifetimes of the L_C_-like domains (Fig. 5).

Stable phase separation has also been reported, employing fluorescence microscopy and AFM, in pulmonary surfactant extract bilayer^24^ and monolayer membranes,^25^ whose behavior was similar when the monolayer was compressed to a pressure of ∼30 mN/m. The monolayer phase behavior is naturally more complex of the two, as it is strongly affected by lateral pressure, as highlighted by AFM experiments.^25^ Importantly, the AFM images measured for monolayers from lung surfactant extracts (compare Fig. 1 with Ref. 25) are very similar to those obtained for our synthetic monolayers. This suggests that our quaternary lipid mixture captures the central features of the phase behavior of the entire complex pulmonary surfactant, suggesting a minor role of surfactant proteins in this respect. Lateral heterogeneity plays a major role in lung mechanics, as pulmonary surfactant needs to possess certain viscoelastic properties in order to decrease the surface tension to low levels upon compression, and at the same time maintain its ability to spread rapidly at the air–water interface, and fold away from the interface to form lipid reservoirs in the aqueous subphase when the lateral pressure exceeds a certain threshold.

The role of DPPC in the pulmonary surfactant has been pictured to reduce surface tension exclusively by forming condensed domains.^34^ This conclusion was drawn from experiments on calf pulmonary surfactant extracts that displayed large flower-shaped DPPC-rich L_C_-like domains. However, these experiments were performed at 293 K, *i*.*e*. 17 K lower than the physiologically relevant body temperature, which can have a major effect on phase behavior near a phase transition. In contrast, our simulations indicate that at room and physiological temperatures, DPPC is only slightly enriched in the L_C_-like domains. Based on such results, we hypothesize that the role of DPPC is to act—together with cholesterol—as a nucleation center for the formation of domains into which lipids with unsaturated chains can also merge. This is further evidenced by our analyses on domain compositions and lifetimes, which suggests that lipids in monolayers have little tendency to undergo phase separation based on the saturation level of their acyl chains (Fig. 4).

In terms of fluidity, POPG has been suggested to be a key player,^68^ in addition to its role in protein–lipid interactions.^18,69^ We observed that bulk L_E_ regions are indeed enriched in POPG. Still, the fluidity is not due to POPG alone, since all the lipid components display fairly similar diffusion coefficients across APL values (Fig. 3). Additionally, we observed that cholesterol has a slight tendency to induce ordered clusters at large APLs, whereas the cholesterols themselves show little tendency to cluster together.

The fact that the spatial heterogeneity was dynamic and dictated by different packing (“physical separation”) instead of the nature of the lipid species (“chemical separation”), suggests that the pulmonary surfactant has specific and collective viscoelastic properties that cannot be derived from the behavior of its components in a straightforward manner. These dynamic properties will be further clarified in our future work. Our simulations also indicate that the behavior of monolayers is in general very similar at room and body temperature. This is not very surprising, as both temperatures fall below the *T*_m_ of DPPC. In addition to these well-balanced viscoelastic properties of the pulmonary surfactant, the observed heterogeneity likely plays a role in regulating the function of surfactant proteins.^18,27^ Computer simulations have become an indispensable tool in biological soft matter research due to their ability to probe small time and length scales.^37^ Traditionally, reproducing surface pressure–area isotherms in classical molecular dynamics simulations has been challenging, and — unlike here — the isotherms in earlier studies were often shifted artificially to match the experimental ones.^70,71^ This signals that in previous simulation studies of pulmonary surfactant layers the descriptions of the physics at the interface has been inadequate due inadequate water models.^44,70^ The simulation approach used in the present work largely overcomes these issues through a combination of recently developed simulation models and simulation parameters.^33,48,49^ The only apparent deviations between the current simulation model and experiment appear at small APLs, where a monolayer collapse plateau is observed in the experimental isotherms at above 45 mN/m and 50 mN/m surface pressures (with areas below 52 Å^2^ and 54 Å^2^) at 298 and 310 K, respectively. This equilibrium collapse pressure is in the ballpark measured for numerous single-component monolayers.^31^ In the simulations at 298 and 310 K, the systems remain stable above the experimental equilibrium collapse pressures of phospholipid monolayers,^31,72^ which likely results from the fact that the simulated monolayer is kinetically trapped in a metastable state and its collapse is limited by the system size and its periodic nature. This limitation has two implications. On the one hand, it hinders studies of the formation of bilayer folds upon high lateral compression, *i*.*e*. during collapse. For such studies, the qualitative picture provided by the MARTINI model^73^ is likely sufficient.^17,29,42^ On the other hand, the atomistic simulations might model well the non-equilibrium situation in the lungs, where monolayer collapse is prevented by rapid compression, and surface pressures can thus reach very high values.^59^ Experimentally, achieving such metastable states requires a specific Langmuir trough or the use of captive bubble surfactometer.^74,75^

One obvious question that arises is whether our AFM observations of sub-micrometer-sized domains are compatible with the nanoscopic and fairly transient domains detected in our simulations. The line tension, *i*.*e*. the penalty of creating a domain boundary, dictates the domain shapes.^24^ The non-circular domain shapes observed by AFM indicate a fairly small line tension, which is also supported by the fact that small and independent domains do not coalesce — as would eventually happen in the case of phase separation. This is in line with the short lifetimes of domains observed in our simulations (Fig. 5C). Based on our simulations, we hypothesize that the L_C_ domains are highly dynamic in monolayers at the air-water interface, but much less dynamic when the samples are transferred onto a mica substrate for AFM measurements. Nevertheless, the heterogeneity observed by AFM is not an artefact of the immobilization, as heterogeneity was also detected in the dilatational modulus of monolayers measured in the Langmuir trough (Fig. 3C). Finally, it is worth noting that the domains visible in our simulations are limited by the simulation box size, despite representing the state-of-the-art in this regard. Our control simulations with more lipids (see Methods) result in similar behavior, thus suggesting that the observed heterogeneity does not arise due to finite-size effects. The surface pressures were also unchanged by the change in simulation box size (Fig. S1). The domain lifetimes extracted from simulations are likely dependent on the domain sizes, and thus also system sizes, since escaping a core of a larger domain takes more time. Unfortunately, extrapolating these lifetimes to microscopic scales is not straightforward.

Our study highlights that, despite the absence of a coexistence plateau in the surface pressure–area isotherm, heterogeneity at the nanometer scale — undetectable by fluorescence or Brewster angle microscopies — should be detected by experimental approaches other than AFM. The surface dilatational rheology measurements using an oscillating barrier readily observed a major change in the dilatational modulus at an APL where the heterogeneities begin to appear (Fig. 3C). Lipid diffusion analyzed from the simulations also revealed a change in the trend at this crossover APL. As monolayer phase transitions are readily detected in experiments that probe lipid diffusion,^61^ such experiments could also detect smaller scale heterogeneity. Finally, lipid tilt measured from our simulations (Fig. 3B) revealed a persistent tilt angle of 25^°^ up to the crossover APL, and this behavior should be captured by either X-ray diffraction^64^ or vibrational sum frequency generation spectroscopy methods.^65^

## Conclusions

Using a combination of Langmuir trough experiments, AFM imaging, and atomistic molecular dynamics simulations, we demonstrated that a synthetic quaternary lipid mixture is able to qualitatively reproduce the key features of the phase behavior of the native pulmonary surfactant extracts. Under a large range of compression levels, thicker L_C_-like domains appear in the otherwise thinner L_E_-phase monolayer. These domains are dynamic and only slightly enriched in DPPC with two saturated chains. We demonstrated that despite there not being a visible phase transition to the L_C_ phase, some monolayer properties change significantly at well–defined values of area per lipid, and this crossover value is consistent between numerous quantities. Moreover, since these properties should be readily measurable using experimental methods, our study also guides experimental work on detecting heterogeneities in biofilms.

Synthetic pulmonary surfactants mimicking the properties of the full functional surfactant are continuously investigated.^76^ A critical aspect is to find a proper lipid composition mixture. Our results highlight how already a few key lipid components of the pulmonary surfactant display small domains, resembling the behavior of surfactant extracts.^25^ The lipid mixture is able to pack in a dynamic manner, thus enabling efficient surface tension reduction while maintaining sufficient fluidity. This behavior might also be crucial for the function of surfactant proteins, as has been investigated in other molecular simulations with multilipid components,^77^ which we will focus on in our future work.

Our approach, combining the CHARMM36 force field with the four-point OPC water model, enables atomistic studies of lipid structures at the air–water interfaces in the complex pulmonary surfactant, allowing for studies of the physiologically important processes in the lung at a detail difficult to achieve experimentally. By integrating experimental data with molecular simulations, we provide, for the first time, a quantitatively accurate, unprecedented picture of the structural and dynamic properties of a realistic model of lung surfactant, under physiologically relevant conditions.

## Acknowledgement

We thank CSC - IT Center for Science (Espoo, Finland) for computing resources. JL and IV thank the Sigrid Juselius Foundation and the Academy of Finland (Centre of Excellence program) for financial support. IV also thanks the HiLIFE (Helsinki Institute of Life Science) Fellow program. RP thanks the Mary and Georg Ehrnrooth Foundation, the Eye and Tissue Bank Foundation, the Evald and Hilda Nissi Foundation and the Biomedicum Helsinki Foundation for financial support. LM acknowledges funding by the Institut national de la santé et de la recherche médicale (INSERM). MJ thanks the Emil Aaltonen Foundation for funding.

