## Supplementary Material for "Understanding the Functional Properties of Lipid Heterogeneity in Pulmonary Surfactant Monolayers at the Atomistic Level"

‡*These authors contributed equally*

⊥*MEMPHYS – Centre for Biomembrane Physics*

### Simulation Details

#### Details on Simulation Models and Methods

Four component lipid monolayers mimicking the native lipid composition of the pulmonary surfactant were simulated with different values for the average area per lipid (APL). A system containing two monolayers with a quaternary mixture of 60 mol-% dipalmitoylphosphatidylcholine (DPPC), 20 mol-% 1-palmitoyl-2-oleoylphosphatidylcholine (POPC), 10 mol-% 1-palmitoyl-2-oleoylphosphatidylglycerol (POPG), and 10 mol-% cholesterol, with a total of 338 lipids (169 per monolayer) with an average APL of 50 Å$^2$, a total of 27 040 water molecules (80 per lipid), and an ion concentration of 0.15 M was generated. The monolayers were separated by a slab of water with air (vacuum) on both sides. The box size in the z direction was set to 22.00 nm to avoid unscreened electrostatic interactions through the vacuum. The system was energy minimized and simulated for 50 ns without restraints with the CHARMM36/OPC4 parameter set at 310 K.

Next, the monolayer structure was either expanded, or compressed during a 10 ns simulation to an average APL value of 100 Å$^2$, or 40 Å$^2$, respectively, using the `MOVINGRESTRAINT` and `CELL` keywords in the PLUMED 2.2 package.[1] From the expansion simulation, frames corresponding to APLs of 52.5, 55, 57.5, 60, 62.5, 65, 67.5, 70, 75, 80, 85, 90, 95, and 100 Å$^2$ were extracted. From the compression simulations, frames corresponding to APLs of 47.5, and 45 were extracted. Independent repetitions at selected APLs of 55, 65, and 75 Å$^2$ were also performed.

These quaternary lipid monolayers were simulated at both 310 K and 298 K for 1.0 µs. The suggested parameter set for CHARMM36 force field combined with the OPC4 water model were used in all monolayer simulations. The simulations were performed in the NVT ensemble and the dispersion correction[2] was applied to both energy and pressure.

We also repeated selected simulations using larger monolayer models. The initial structures for the large quaternary lipid monolayers were taken as the final configurations of selected smaller monolayer simulations described above. The structures were multiplied using the g_genconf tool, to generate a system with a total of 1352 lipids (676 per monolayer) and a total of 108160 water molecules, and a system with a total of 3042 lipids (1521 per monolayer) and a total of 243360 water molecules. All systems had an ion concentration of 0.15 M. The simulation parameters were identical to the ones used with the small monolayers. The 1352 lipid systems with APLs of 50.0 and 55.0 Å$^2$ at 298 K and 310 K, respectively, were simulated for 1 µs both. The 3042 lipid system with APL of 55.0 Å$^2$ at 310 K was simulated for 500 ns.

#### Calculation of $\gamma_0$

The pure air–water interface was simulated using the four-point OPC[3] (OPC4) water model to evaluate the interfacial surface tension values ($\gamma_0$) used in the calculation of the surface pressure of the monolayers. The systems contained 27040 water molecules for the smaller, and 108160 water molecules for the larger systems, with box dimensions of $9.19^2 \times 22.00$ nm$^3$, and $19.28^2 \times 22.00$ nm$^3$, respectively, corresponding to the average sizes of the monolayer simulations described above. The OPC4 water model with the suggested simulation parameters for the CHARMM36 force field with GROMACS was used (see Ref. 4 for details). The systems containing 27040 water molecules were simulated at 298 K and 310 K for 30 ns, and the systems with 108160 water molecules were simulated at 298 K and 310 K for 10 ns. The simulations were performed in the NVT ensemble and the dispersion correction[2] was applied to both energy and pressure. The first 10 ns of the simulations were omitted from the analysis of the smaller water systems and the first 1 ns were omitted from the analysis of the larger water systems. All simulations were run with GROMACS 5.1.x.[5] The surface tension of water ($\gamma_0$) was extracted using the g_energy tool.

### Supplementary Experimental Results

#### Pressure–Area Isotherms

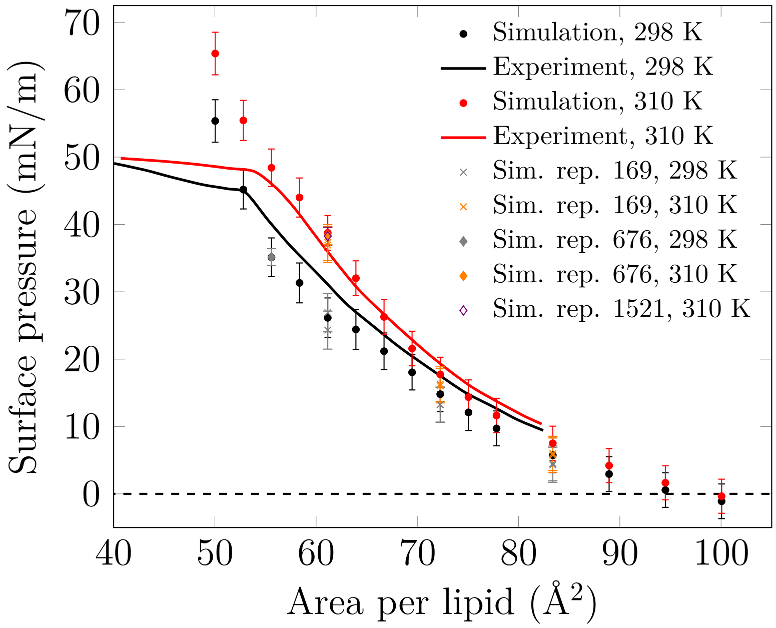


Figure S1: Surface pressure–Area isotherms at 298 K and 310 K. Due to the periodic boundary conditions and the finite sizes of the systems, the monolayers cannot collapse in the simulated time scale. Therefore, the simulations overestimate the surface pressures of the quaternary monolayers with areas below 53 and 56 Å$^2$ at 298 K and 310 K, respectively, where the monolayers are mostly in the $L_c$ phase and in a metastable state. Replicas were performed for certain APL values to check the consistency of the calculated surface pressure values. Moreover, additional simulations with larger monolayers (676 or 1521 lipids per monolayer) were performed to evaluate the finite-size effects on the calculated surface pressure values.

#### AFM Imaging

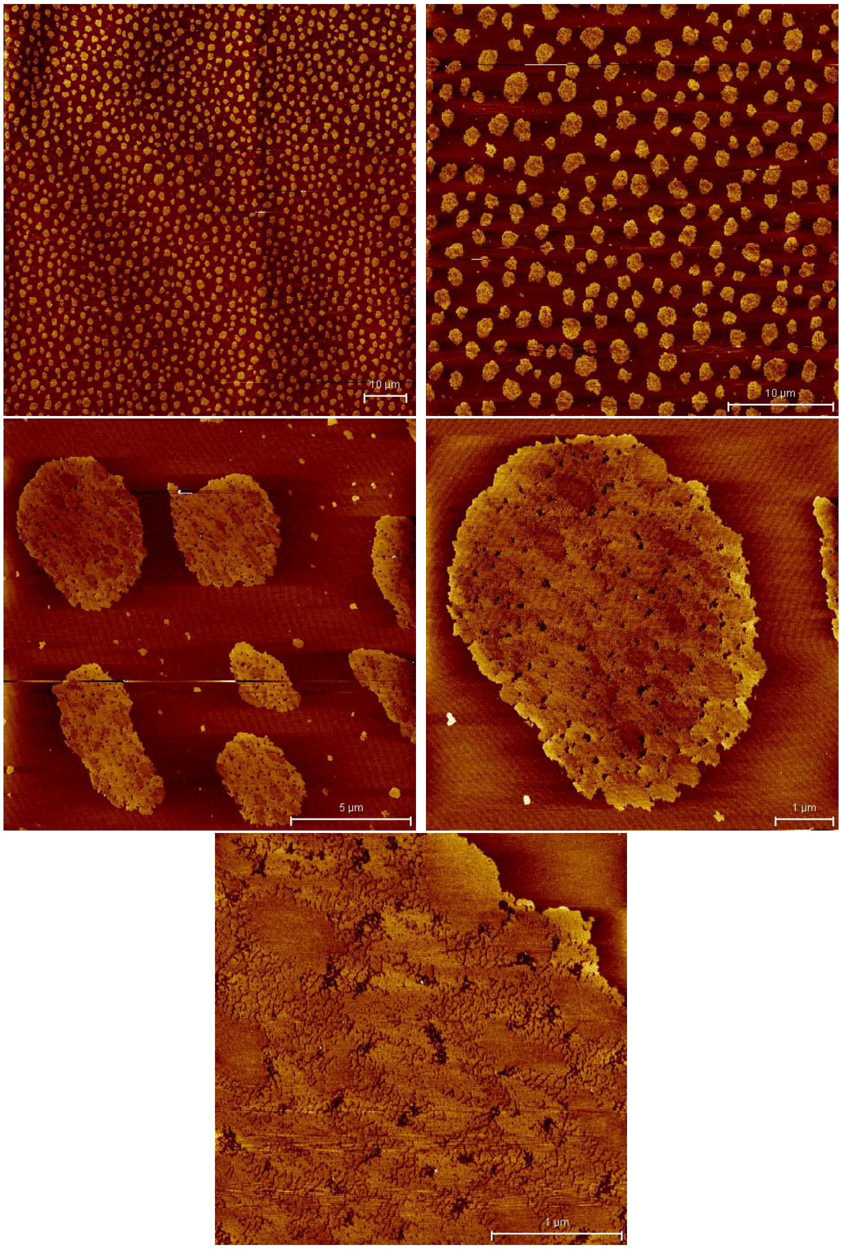


Figure S2: AFM images at 25 mN/m.

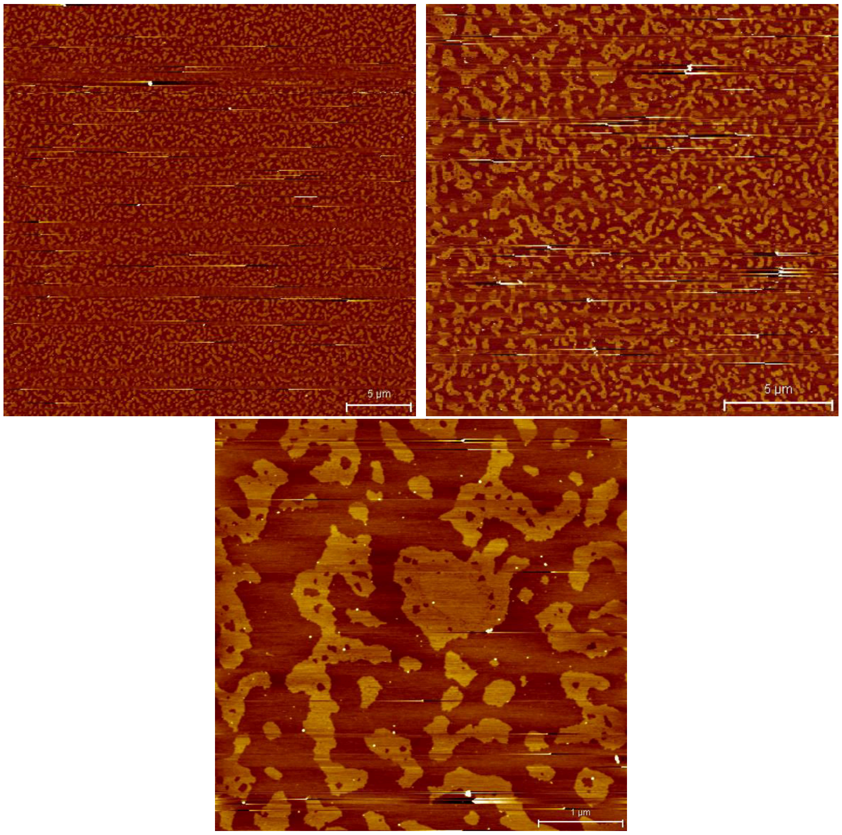


Figure S3: AFM images at 35 mN/m.

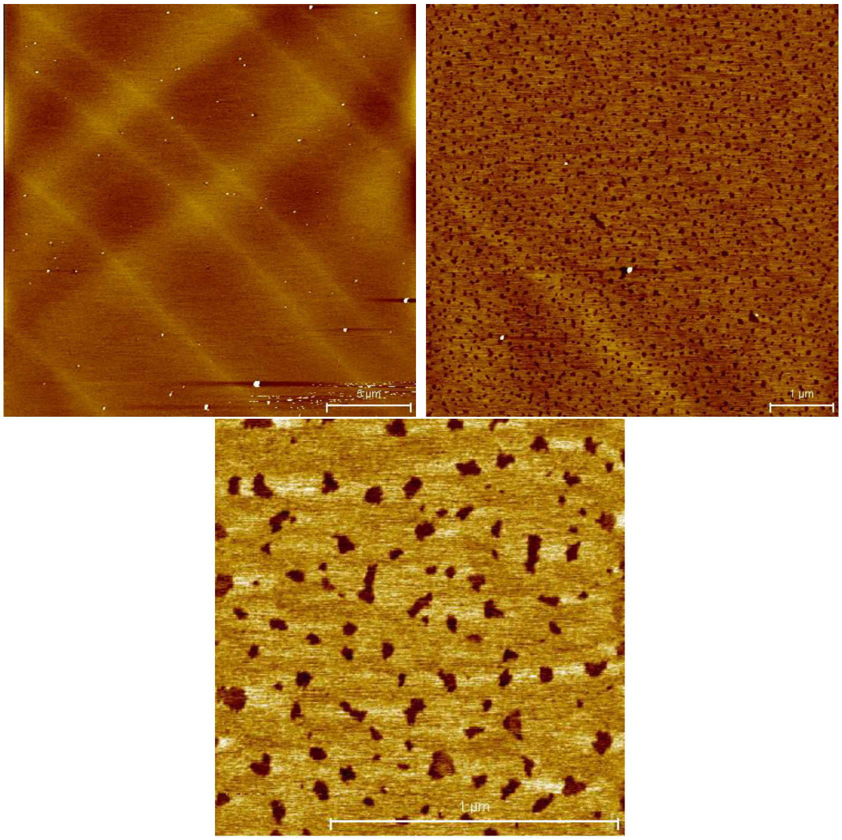


Figure S4: AFM images at 45 mN/m.

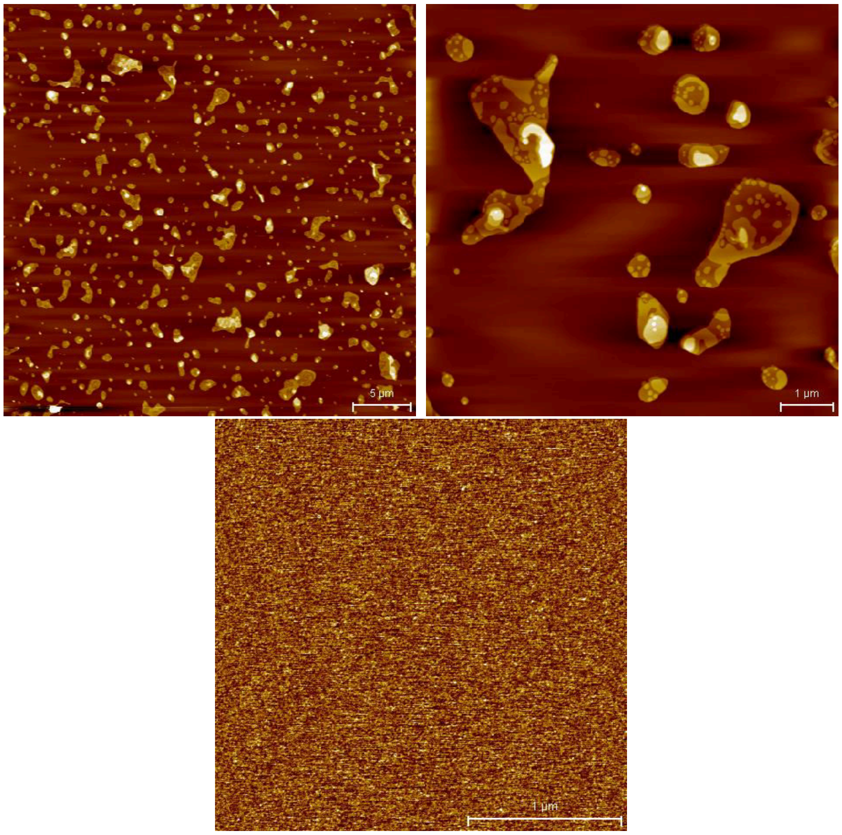


Figure S5: AFM images at 55 mN/m.

#### Height Profile Studies

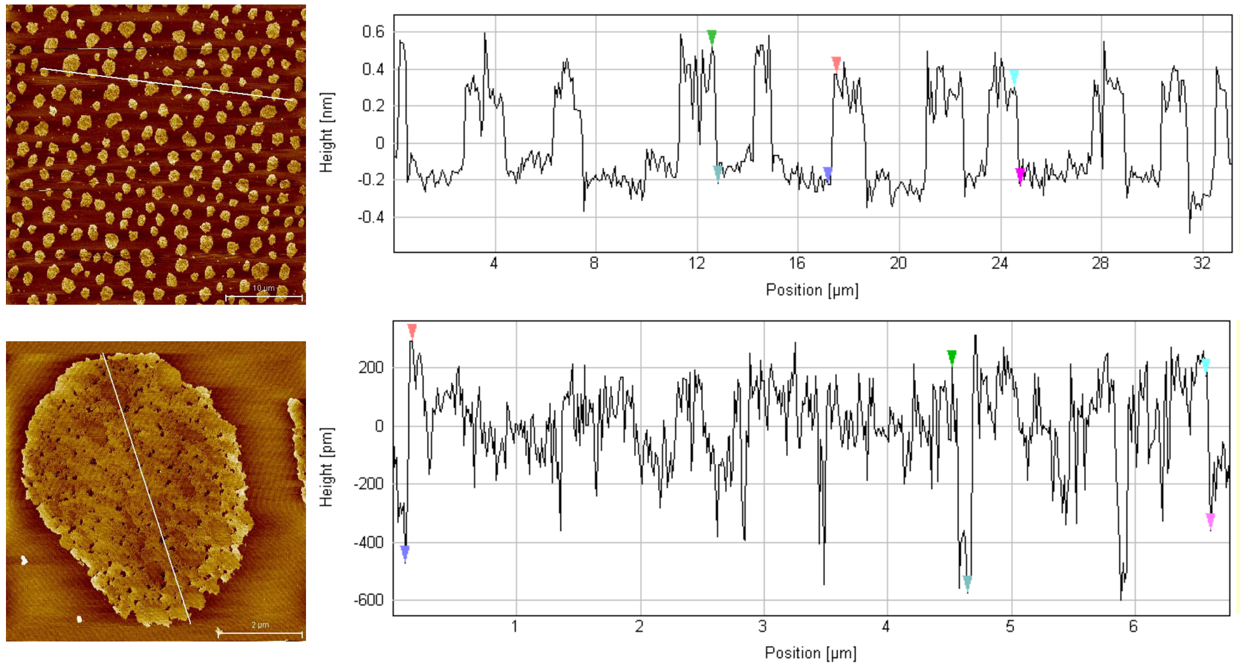


Figure S6: Heigth profile study at 25 mN/m.

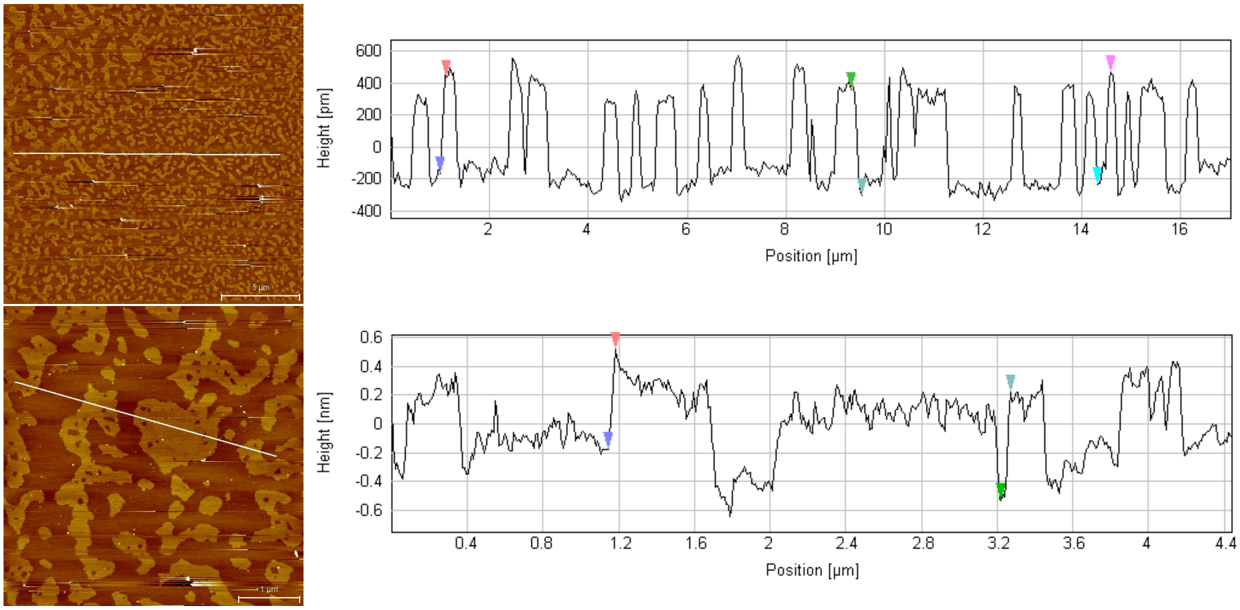


Figure S7: Heigth profile study at 35 mN/m.

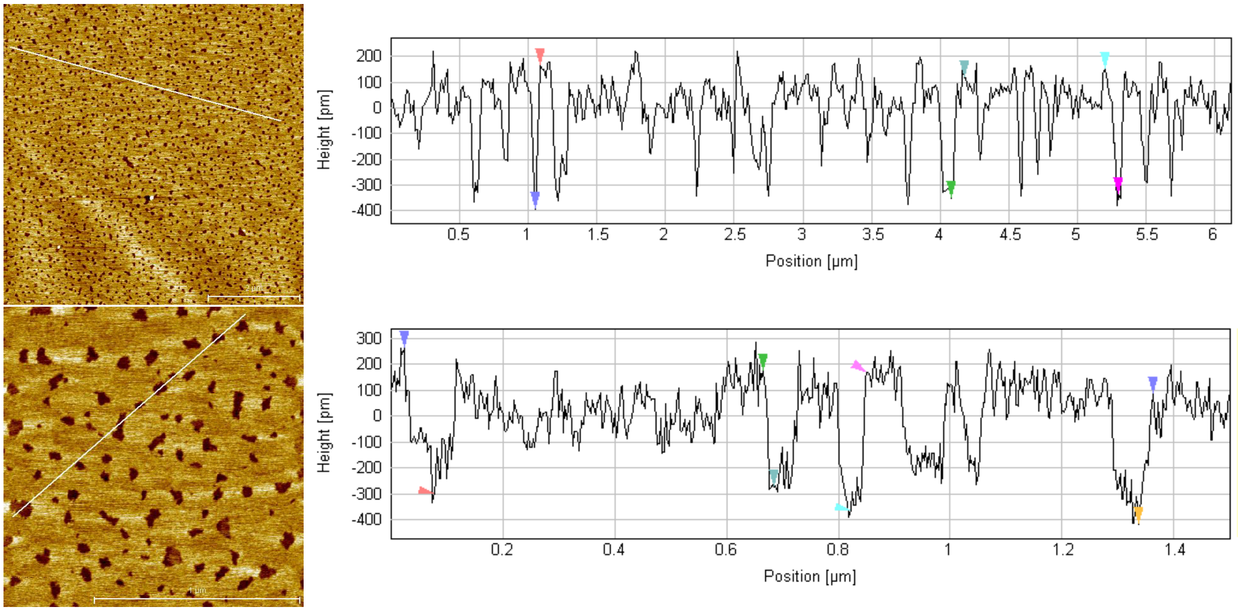


Figure S8: Heigth profile study at 45 mN/m.

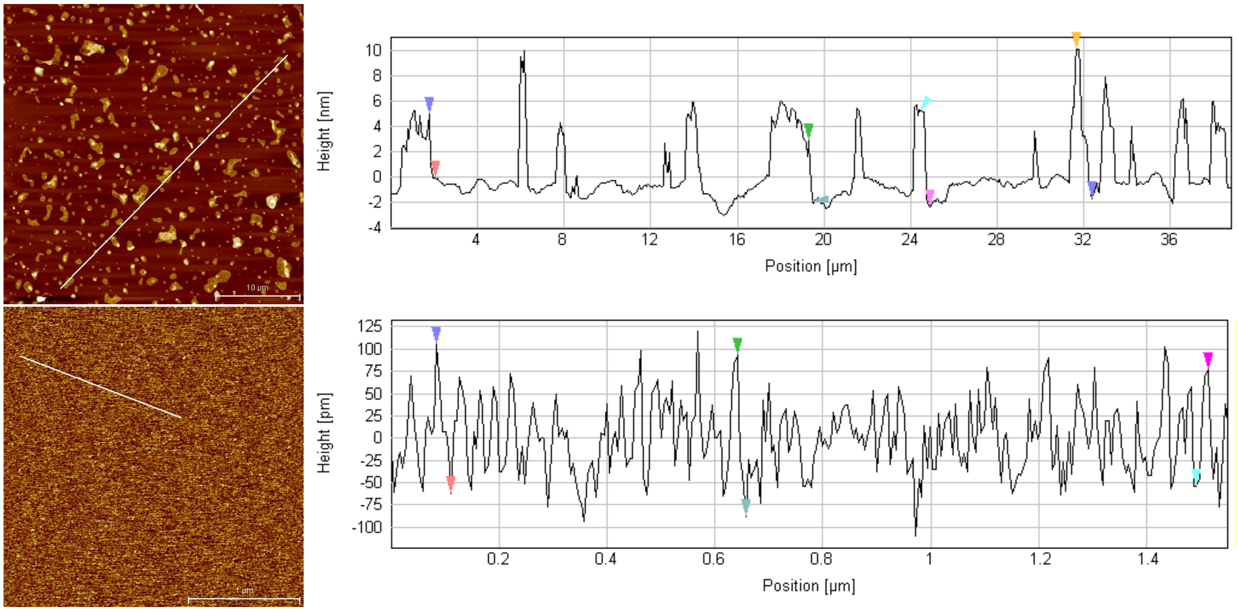


Figure S9: Heigth profile study at 55 mN/m.

### Supplementary Simulation Results

#### Snapshots of the Lipid Monolayer Simulations

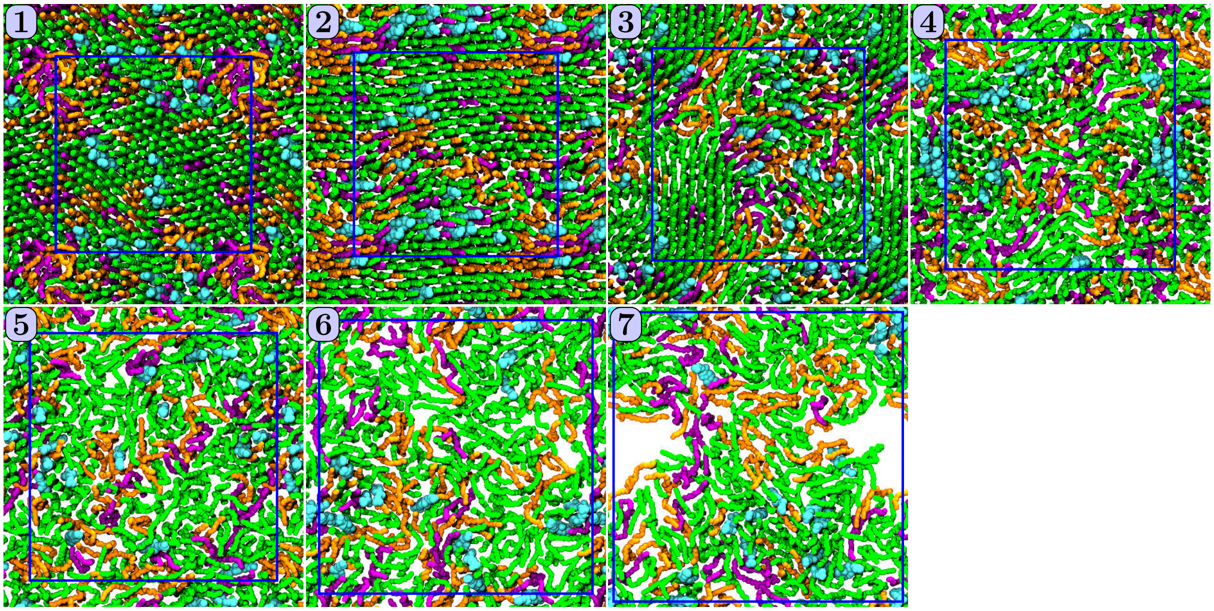


Figure S10: Snapshots of the lipid monolayers at 298 K. Top row, from left to right: 45 Å$^2$, 50 Å$^2$, 55 Å$^2$, and 65 Å$^2$. Bottom row: 75 Å$^2$, 90 Å$^2$, and 100 Å$^2$. Blue lines highlight the simulation cell.

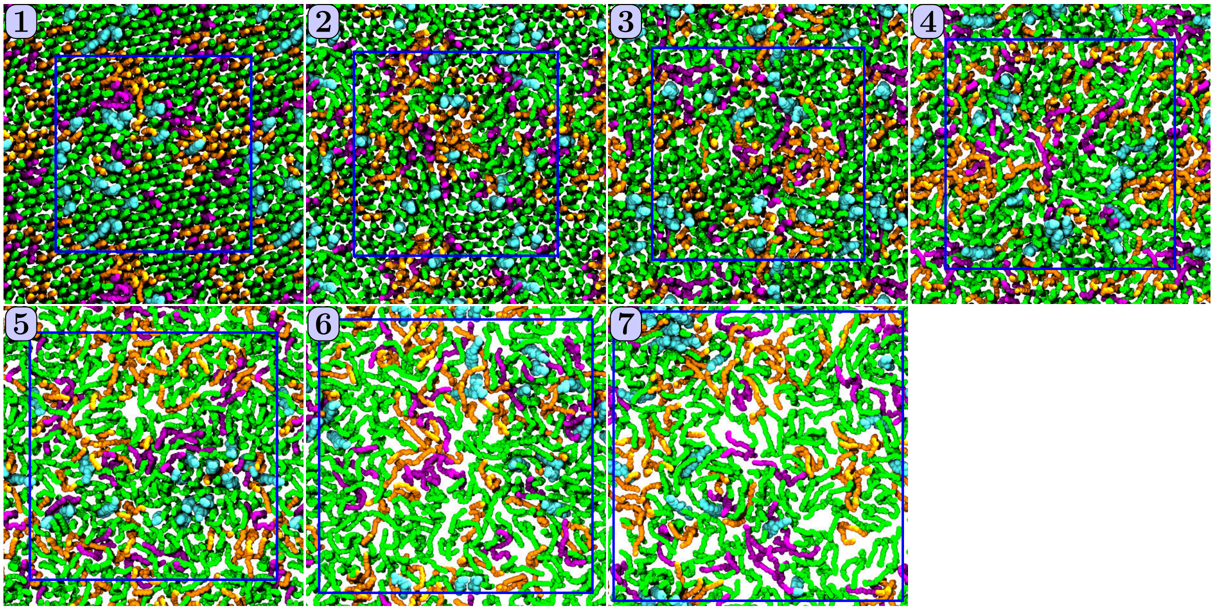


Figure S11: Snapshots of the lipid monolayers at 310 K. Top row, from left to right: 45 Å$^2$, 50 Å$^2$, 55 Å$^2$, and 65 Å$^2$. Bottom row: 75 Å$^2$, 90 Å$^2$, and 100 Å$^2$. Blue lines highlight the simulation cell.

#### Tilt Angle Distributions

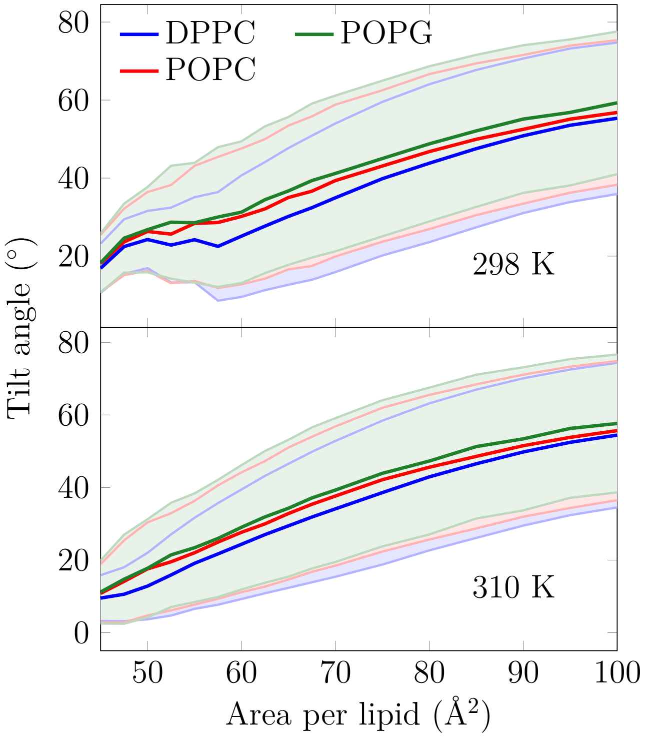


Figure S12: Tilt angle distributions of lipid chains. The shaded areas, bordered by dim lines, show the standard deviation.

#### Cholesterol clustering

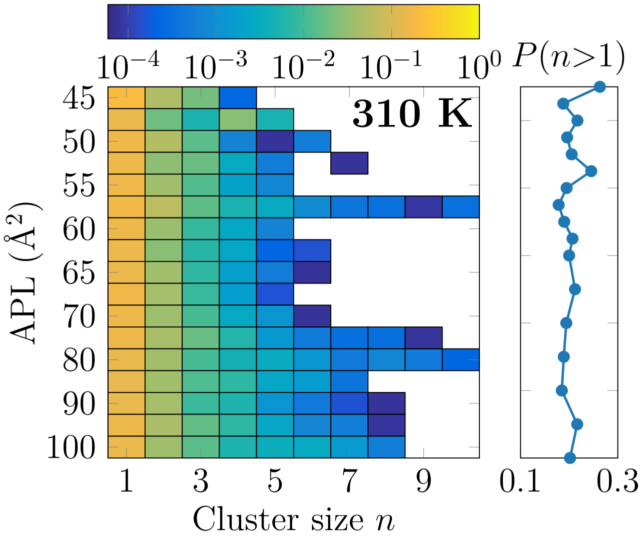


Figure S13: Left: 2-dimensional probability distributions of cholesterol molecules residing in a cluster with at least one other cholesterol at all simulated APLs at 310 K. Note that the color bar is in logarithmic scale. Right: The fraction of cholesterol molecules in clusters with size larger than one.

#### Availability of Simulation Data

Links to Zenodo repositories for simulation data with descriptions and links will be added
